# Function-preserving CD8⁺ T Cell Immunogenicity minimization through protein design

**DOI:** 10.64898/2026.10.24.761497

**Authors:** Bingxu Liu, Tianyang Mao, Max H Witwer, Oliver Morell, Nir Hacohen, David Baker

**Affiliations:** Department of Biochemistry, University of Washington, Seattle, WA 98195, USA; Institute for Protein Design, University of Washington, Seattle, WA 98195, USA; Broad Institute of MIT and Harvard, Cambridge, MA 02142, USA; Massachusetts General Hospital Cancer Center, Boston, MA 02129, USA; Howard Hughes Medical Institute, University of Washington, Seattle, WA 98195, USA; Department of Biotechnology and Biomedicine, Technical University of Denmark, Kongens Lyngby, 9 Denmark

## Abstract

Rapid advances in protein design enable new approaches in adoptive cell therapy, but foreign proteins can elicit cytotoxic T cell responses, and hence there is considerable need for approaches which reduce immunogenicity without compromising protein function. Here, we develop a computational framework integrating structure-guided sequence design with MHC class I epitope prediction to efficiently tune protein immunogenicity while preserving function. Using a de novo-designed luciferase as a model, we demonstrate epitope elimination while preserving function across 5 common mouse alleles and extend our computational optimization to 25 common human MHC-I alleles. In mice immunized with mRNA-lipid nanoparticles, variants designed to have few MHC-I epitopes elicited little detectable CD8⁺ T cell response, whereas variants designed to include multiple epitopes induced robust responses. Together, these results demonstrate that the CD8⁺ T cell immunogenicity of designed proteins can be tuned while preserving function, opening the door to incorporation of designed proteins in a next generation of adoptive cell therapies.

## Main text

Designed proteins have been widely adopted for cell recognition^1–3^, cell signaling^4–7^, and show great promise in fields like adoptive cell therapies^2–4^. However, protein immunogenicity presents a major barrier to clinical application^8–11^. Intracellular proteins are continuously processed into short peptides that can be presented by major histocompatibility complex class I (MHC-I) molecules, potentially eliciting cytotoxic T cell responses and elimination of cells expressing the therapeutic protein. Foreign proteins (such as resistance genes^8,11^) or even modified human proteins^10^ (that have not been tolerized in recipients) expressed in cells can lead to cytolytic T cell induction and elimination of the cells expressing these genes. Removing immunogenic epitopes is challenging as extensive sequence modifications may compromise protein folding and function. Thus, a central challenge is to redesign proteins to minimize immunogenicity while preserving their biological activity.

Advances in generative protein design could provide a potential solution to this challenge. Structure-conditioned sequence design tools, such as ProteinMPNN^12^, enable extensive exploration of sequence space while maintaining the desired protein structures that are essential for their functions. In parallel, MHC-I presentation predictors, such as NetMHCpan4.1^13^, enable computational identification of potentially immunogenic peptides. Previous studies have demonstrated ProteinMPNN-based sequence redesign to reduce predicated MHC-I epitopes^14–17^ but whether such design methods can completely remove CD8⁺ T-cell immunogenicity responses validated *in vivo* and be expanded to dozens of alleles remain insufficiently explored. We set out to develop a computational deimmunization method that preserves protein function while eliminating epitopes across multiple MHC-I alleles, and investigate the immunogenicity of the designs in animal models.

## Functional proteins can be designed with high or low predicted MHC-I epitope burdens

To reduce protein immunogenicity while preserving protein function, we developed an iterative optimization strategy integrating ProteinMPNN and NetMHCpan4.1, called MINT (MHC-I Immunogenicity Tuning; Figure 1A). Starting from an input 3D structure, we use ProteinMPNN to generate a diverse set of output sequences, identify class I epitopes with NetMHCpan4.1 over an input selected set of MHC-I alleles, and evaluate folding to the input structure using structure prediction methods. If the set of MHC alleles is small, typically at least one of the generated sequences will both lack epitopes and be predicted to fold to the target structure, and be suitable for experimental characterization. To design proteins that avoid presentation by the larger set of human HLA alleles from the human population, this zero shot procedure is typically insufficient, and in this case we carry out iterative rounds of mutation and selection while retaining pool diversity, selecting at each stage diverse variants with low predicted immunogenicity (Figure 1A, see Methods). For the mutation step, both MPNN-guided multiple-site sequence redesign and saturation point mutation were applied to the predicted epitopes (Figure 1A and methods). For immunogenicity prediction, NetMHC computation is a major time cost; to improve search efficiency, we cached NetMHCpan predictions for peptide-allele pairs so they could be reused across candidates and rounds.

**Figure 1.**
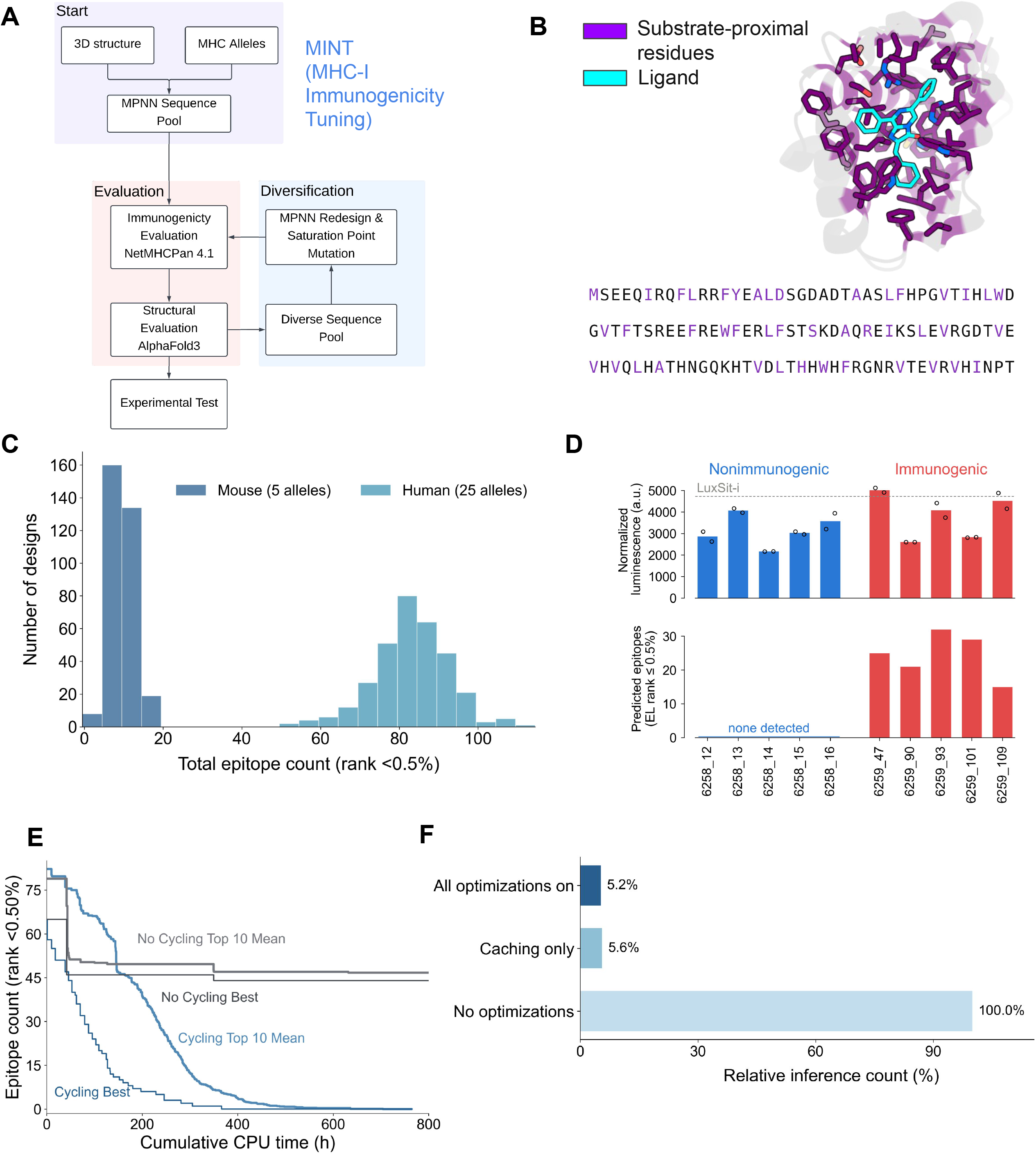
Structure-guided protein design enables modulation of predicted MHC-I epitope burden while preserving luciferase activity. (A) Schematic of MINT deimmunization algorithm. (B) Predicted structure and amino acid sequence of LuxSit-i. The substrate Diphenylterazine (DTZ) is shown in cyan, and residues constrained during sequence redesign are highlighted in purple. (C) Distribution of the number of strong (EL rank <0.5%) MHC-I epitopes for 25 human MHC-I alleles (cyan) and 5 mouse MHC-I alleles (navy) from a single pass. (D) Top, Normalized luminescence of ten designed LuxSit-i variants. Each dot is a technical repeat. Bottom, Number of predicted strong MHC-I-epitopes (EL rank <0.5%) across five mouse MHC-I alleles for ten LuxSit-i variants. (E) Cumulative CPU hours and predicted MHC-I epitope reduction for MINT with cycling (blue) and without cycling (gray) evaluated across 25 human alleles. Without cycling 500,000 designs were generated; with cycling, 20 design campaigns were run in parallel. Dark lines show the best result discovered by each method. For no cycling, the light lines show the average epitope count among the 10 best individual designs. For cycling, they show the epitope count of the best design from each of the 10 best-performing campaigns. Predicted epitopes are defined by EL rank <0.5%. (F) Ablation analysis showing number of MixMHCpred 4.1 evaluations required at different levels of MINT optimization to assess the same 600 candidate sequences. Values are expressed as a percentage of no optimizations workload: All optimizations on (top bar), Caching only (middle bar), and no optimizations (bottom bar).

To test the performance of MINT, we chose a de novo designed luciferase (LuxSit-i^18^ (Figure 1B) as a model system. Luciferase is widely used in biological imaging, and activity can be readily measured, enabling rapid assessment of retention of protein function. We used MINT to redesign LuxSit-i for different class I allele sets while constraining 43 substrate-proximal residues to preserve enzymatic activity (Figure 1B). The resulting sequences exhibited substantial diversity in amino acid composition (Figure S1A), demonstrating that our method does not converge to a narrow set of solutions.

When tasked with removing epitopes for 5 common mouse alleles, MINT in a single pass identified LuxSit-i variants predicted to have low immunogenicity, alongside variants predicted to have high immunogenicity (Figures 1C, 1D, and S1B). The low and high predicted immunogenicity subsets retained comparable luciferase activity (Figure 1D), indicating separation of immunogenicity from protein function.

Human populations have a much larger set of alleles, and removing epitopes for larger numbers of alleles simultaneously is a more challenging optimization challenge (Figure S1B). When tasked with removing epitopes for 25 human alleles, MINT required multiple cycles of mutation and selection to fully deimmunize Luxsit-i (Figure 1C and 1E). Caching to avoid repetitive NetMHC calls considerably increased efficiency (Figure 1F), making it feasible to deimmunize completely.

## Luciferase can be designed to elicit minimal or strong CD8⁺ T cell responses in vivo

To determine whether the computationally designed differences in MHC-I epitope burden translate into differences in T cell immunogenicity, we selected the five predicted high-epitope and five predicted no-epitope luciferase variants with comparable enzymatic activity for immunogenicity assessment in vivo (Figure 1D). We employed mRNA-lipid nanoparticles (LNP) as the vaccine platform, which efficiently induces antigen-specific T cell responses^19–21^, providing a sensitive method for assessing the immunogenic potential of the designed proteins.

To compare responses to multiple luciferase variants within a shared immunization context, we constructed five synthetic mRNAs encoding concatenated luciferase proteins, each containing all five predicated no-epitope variants together with one predicted high-epitope variant connected by flexible linkers (denoted as the “5+1” fusion; Figure 2A). Mice were immunized with mRNA-LNPs encoding each fusion using a prime-boost regimen, and splenocytes were collected seven days after boosting to evaluate antigen-specific T cell responses (Figure 2B).

**Figure 2.**
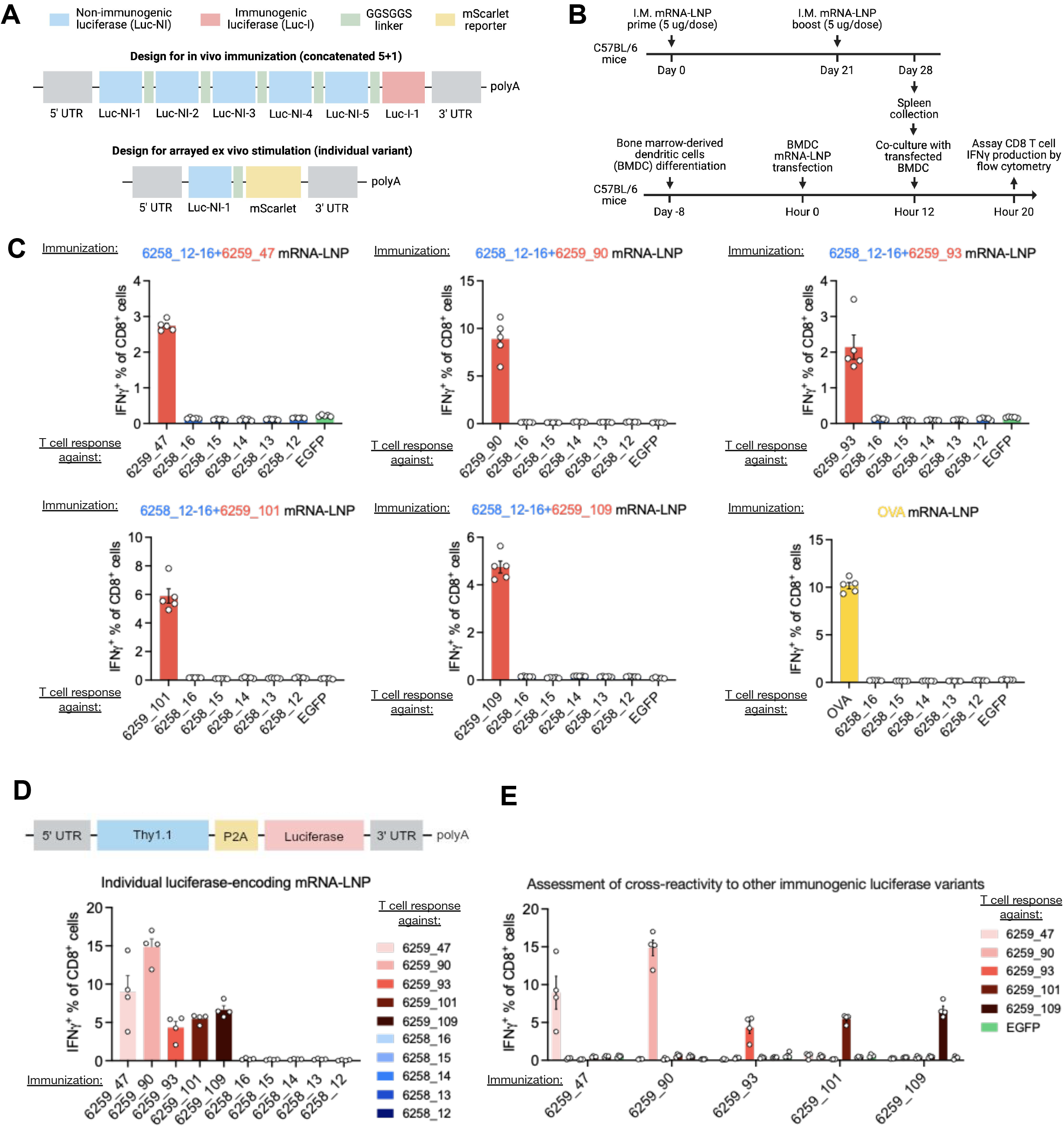
Designed luciferase variants elicit distinct CD8⁺ T cell responses in vivo. (A) Schematic of the 5+1 fusion constructs containing five predicted low-immunogenicity variants and one predicted immunogenic LuxSit-i variant. (B) Schematic of the mRNA-LNP immunization regimen and subsequent assessment of antigen-specific CD8⁺ T cell responses. (C) Frequencies of IFN-γ⁺ CD8⁺ T cells responding to individual LuxSit-i variants following immunization with 5+1 fusion constructs. (D) Top: schematic of the individual-variant immunization experiment. Bottom: frequencies of IFN-γ⁺ CD8⁺ T cells responding to individual LuxSit-i variants following immunization. (E) Cross-reactivity of CD8⁺ T cell responses elicited by individual immunogenic LuxSit-i variants against other immunogenic variants.

To measure T cell responses against individual luciferase variants, we established an ex vivo antigen-presentation assay using bone marrow-derived dendritic cells (BMDCs). BMDCs were transfected with synthetic mRNAs encoding individual luciferase variants together with an mScarlet fluorescent reporter (Figure 2A). After confirming successful transfection based on reporter fluorescence (Figure S2A), we co-cultured the BMDCs with splenocytes from immunized mice and measured antigen-specific CD8⁺ T cell responses (Figure 2B). We observed robust responses to predicted high-epitope luciferase variants, characterized by IFN-γ, TNF-α, and IL-2 production, whereas predicted no-epitope variants elicited minimal detectable responses within the same immunization context (Figures 2C and S2B–D).

Although these results demonstrate differential T cell recognition of designed luciferases within the same immunization context, competition between epitopes in the 5+1 fusion could potentially suppress responses to predicted no-epitope variants^22^. To distinguish intrinsic differences in immunogenic potential from antigenic competition, we generated ten additional mRNA-LNP immunogens, each encoding an individual luciferase variant from the five predicted high-epitope and five predicted no-epitope designs (Figures 2D). Reporter expression confirmed successful transfection of BMDCs with constructs encoding the individual variants (Figure S2E).

Consistent with the multivalent immunization results, individual predicted high-epitope variants elicited robust CD8⁺ T cell responses, whereas predicted no-epitope variants induced minimal detectable responses (Figures 2D). Furthermore, T cells elicited by individual predicted high-epitope designs preferentially recognized their corresponding luciferase variants, with limited cross-reactivity toward other designs (Figure 2E), highlighting the designed sequence diversity. Together, these results demonstrate that protein sequence redesign can substantially alter antigen-specific CD8⁺ T cell responses.

## Discussion

We demonstrate that CD8⁺ T cell immunogenicity can be systematically modulated through protein sequence redesign while preserving enzymatic activity. By generating functional luciferase variants with contrasting immune responses, we establish an experimental connection between computational MHC-I epitope optimization and antigen-specific T cell recognition in vivo. These findings expand the scope of protein design beyond conventional objectives such as stability, structure, and biochemical function, establishing immunogenicity as an additional property that can be rationally designed.

An advantage of our approach is the modularity in defining immunogenicity as a sequence design objective. By decoupling structure-guided sequence generation from immunogenicity evaluation, our approach allows different prediction models, HLA allele combinations, and optimization objectives to be incorporated without retraining the underlying protein design model. This flexibility is particularly valuable as immunogenicity prediction methods continue to improve. While our experimental validation focused on MHC-I–restricted CD8⁺ T cell responses, the same framework could incorporate MHC-II epitope predictors to guide the engineering of CD4⁺ T cell recognition. This could support future efforts in antibody-response deimmunization as MHC-II prediction models improve in their ability to capture in vivo immunogenicity^23,24^. Protein design can now generate potential therapeutic candidates ranging from highly specific binding proteins to enzymes, and approaches such as MINT should be broadly applicable for reducing their immunogenicity. Conversely, for inducing T cell responses against antigens introduced into tumors, MINT could be useful for maximizing epitope numbers for systemic anti-immunity through antigen spreading^25,26^. Together, these advances provide a modular foundation for designing proteins with tailored immunogenicity while preserving biological function.

## Acknowledgements

This research was supported by Cancer Grand Challenges team MATCHMAKERS funded by Cancer Research UK CGCATF-2023/100008 (B.L., D.B.); the National Cancer Institute 1OT2CA297288-01 (D.B.); and The Mark Foundation for Cancer Research (N.H., D.B.); Department of Defense, Defense Threat Reduction Agency grant HDTRA (B.L.); Microsoft Protein Prediction (B.L., D.B.); Jane Coffin Childs Memorial Fund for Medical Research (T.M.), D.B. is Investigator of the Howard Hughes Medical Institute. We thank Sam Pellock for helping with LuxSit-i structure and function analysis, Stephanie Dougan, Michael Dougan, Tavus Atajanova, Qiang Li for in vivo experiment discussion and exploration, Morten Nielsen for discussion on NetMHCpan4.1 models, Michael Birnbaum for suggesting the human alleles to prioritize deimmunize against, Timothy Jenkins, Kejia Wu and Jeremiah Sims for discussions on algorithmic improvements, GeneFrontier Corporation for kind donation of Cell free expression reagents,

## Method

### Luciferase activity measurement

LuxSit-i variants were ordered as linear DNA constructs encoding fusion proteins with mScarlet-I3. The DNA constructs were expressed using an *in vitro* transcription and translation (IVTT) system (gift from Gene Frontier, Purefrex 2.1) at 37°C for 2–4 h. Following expression, 2 uL IVTT reaction products were mixed with the luciferase substrate 30 uL 50 uM Diphenylterazine (DTZ) (Medexpress) in a 96-well white plate. Fluorescence and luminescence were measured using BioTek Synergy Neo2. Luciferase activity of variants was normalized by the mScarlet fluorescence signal for each sample.

### mRNA synthesis

LuxSit-i variant-encoding sequences were codon optimized, synthesized, and cloned into a T7 in vitro transcription vector (Takara). In vitro transcription was performed in 20 µL reactions containing 1 µg linearized template, 1× T7 Reaction Buffer, 10 mM each of ATP, GTP and CTP, 10 mM N1-methylpseudouridine-5′-triphosphate (m1Ψ, TriLink BioTechnologies) in place of UTP, 8 mM CleanCap AG (TriLink BioTechnologies) and T7 RNA polymerase mix (NEB)^27,28^. Reactions were incubated at 37 °C for 2 h, then treated with RNase-free DNase I (NEB) for 15 min at 37 °C to remove template DNA. Transcripts were purified on PureLink RNA Mini columns (Thermo Fisher Scientific), quantified by NanoDrop and stored at −80 °C until formulation.

### mRNA-LNP formulation and characterization

Lipid nanoparticles were formulated with a clinically validated ionizable lipid composition with SM-102, DSPC, cholesterol and DMG-PEG2000 mixed at 50 : 10 : 38.5 : 1.5 mol%^29,30^. The aqueous and organic phases were combined at a 2:1 (v/v) ratio on a Spark microfluidic formulation instrument, with an ionizable lipid nitrogen to RNA phosphate (N:P) molar ratio of 6:1. Ethanol was removed using 10 kDa MWCO spin concentrators (2,500 × g, 4 °C, 45 min). Encapsulation efficiency was determined with the Quant-iT RiboGreen RNA Assay Kit (Thermo Fisher Scientific) by comparing RNA signal in TE buffer with signal after lysis in 1% Triton X-100, against an RNA standard curve.

### Mice and immunization

C57BL/6N mice (Charles River) were housed under specific pathogen-free conditions. All procedures were approved by the Institutional Animal Care and Use Committee. mRNA-LNPs were freshly formulated and diluted in sterile 1× PBS on the day of injection and delivered intramuscularly for a total of 5 µg mRNA per mouse per dose. Mice received a prime on day 0 and a boost on day 21. Spleens were harvested on day 28, seven days after boost, into cRPMI and processed immediately to single-cell suspensions for ex vivo restimulation assays.

### Ex vivo CD8 T cell restimulation

Bone marrow was flushed from the femurs and tibiae of CD45.1+ mice (BoyJ; JAX) and cultured at 2 × 10⁶ cells/mL in cRPMI supplemented with 20 ng/mL GM-CSF, 50 ng/mL human Flt3L and 50 µM 2-mercaptoethanol. Eight days post in vitro differentiation, BMDCs were harvested and transfected with mRNA-LNP encoding de novo designed luciferases for 12 h at 37 °C. Splenocytes from immunized mice were added to transfected BMDCs at 5:1 splenocyte-to-BMDC ratio per well containing a Protein Transport Inhibitor Cocktail (Invitrogen) and co-cultured for 8 h at 37 °C. Following co-culture, cells were stained with a Zombie NIR Fixable Viability dye (1:500, 15 min, room temperature), blocked with anti-CD16/32 (1:200, 15 min, room temperature) and surface stained with CD8a-Spark PLUS UV395 (BioLegend, Clone 53-6.7), CD90.2-BV421 (BioLegend, Clone 30-H12), CD44-BV605 (BioLegend, Clone IM7), CD4-BV785 (BioLegend, Clone GK1.5), and CD45.1-FITC (BioLegend, Clone A20) (1:200, 30 min, 4 °C). Cells were then fixed in 4% paraformaldehyde (30 min, 4 °C), permeabilized for 10 min at room temperature using the Foxp3/Transcription Factor Staining Buffer Set (Invitrogen) and stained intracellularly with IFN-γ-PE (BioLegend, Clone XMG1.2), TNF-α-PE-Cy7 (BioLegend, Clone MP6-XT22) and IL-2-APC (BioLegend, Clone JES6-5H4) (1:200, 30 min, room temperature). Samples were acquired on a CytoFLEX LX cytometer (Beckman Coulter). Antigen-specific responses were quantified as the frequency of cytokine-producing cells among live CD90.2⁺CD8⁺ T cells.

### Starting structures and design constraints

LuxSit-i comprises 117 amino acid residues. To preserve catalytic activity, the 43 residues closest to the substrate (37% of the protein) were fixed, leaving 74 positions available for mutational design to modulate predicted HLA class-I presentation. Fixed positions retain their amino acid identities across design rounds, but were not considered to be fixed coordinates during structure prediction. During redesign, the mutable positions were permitted to vary, while the introduction of new cysteine residues was prohibited. During optimization, the initial seed backbone was retained and used as a structural reference against which all redesigned and folded sequences were compared.

### MINT Design Creation without Cycling

To create the deimmunized variants vanilla ProteinMPNN was run for 40,000 backbone redesigns and those having zero peptides below EL-Rank 0.5% were folded with AF3.

To create the immunized variants vanilla ProteinMPNN was run at a sampling temperature of 0.25 for three iterative rounds of redesign, improving the mean Cα pLDDT of the starting design from 82.8 to yield a diverse set of 20 sequences with above 95 pLDDT and aligned Cα RMSD ≤1.0.

The functional designs were then generated using immune guided proteinMPNN sequence design for 14 rounds. Each round began with a step that redesigned the sequence completely driving it away from the parent sequence by down-weighting the probability of the parent residue at each mutable position; this was followed by a second pass incorporating predicted HLA class-I presentation during sequence selection. At each mutable position, all 20 amino acid alternatives were evaluated. For each substitution, all overlapping 9-11-mer scanning windows were evaluated using NetMHCpan 4.1. EL percentile rank was converted into a continuous penalty:

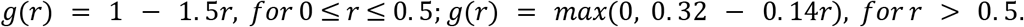

Where r is the predicted EL percentile rank expressed in percent units (e.g. r=2.0 for a 2% rank).

All penalties were summed across the contributing windows and alleles to score a given point mutation. The scores were then scaled by 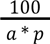 where a is the number of alleles considered and p is the number of peptide lengths considered. The normalized score was then mean-centered across the 20 candidate residues at that position to obtain *I_i_*(*a*). The weighting term on a given amino acid mutation is:

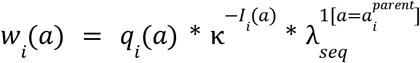

where w is the weight, *q* is the output probability distribution of ProteinMPNN, κ is the immune weighting base and λ is the sequence divergence term. The weighting term is then converted into a probability via temperature guided renormalization:

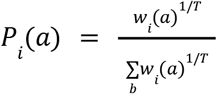

For the aforementioned fourteen rounds, T=0.3, λ=0.1 and κ=0.95.

This inference practice for designing more immunogenic variants is optional for generation of sequences, and the simpler method for generating deimmunized variants can be adapted to find immunogenic sequences.

### MINT Design Creation with Cycling

#### 1. Design Library

A library of 20 sequence-divergent Luxsit-I variants was generated by altering mutable positions with vanilla ProteinMPNN and evaluating the resulting sequences with AF3. Variants were retained if they achieved a mean Cα pLDDT score of at least 95 and an aligned Cα RMSD of no more than 1.0 Å relative to the starting reference structure. This approach produced backbones with modest conformational differences, allowing us to test whether 20 distinct sequence lineages with subtly different predicted backbone structures converged on similar predicted HLA class-I presentation profiles.

#### 2. Optimization Objective

To score a protein sequence, all peptide windows of length 9-11 residues were enumerated and scored using NetMHCpan 4.1 EL percent ranks against a panel of 25 HLA class-I alleles (Figure 1C). Cumulative counts were recorded below 2% and 0.5% rank, with the latter included in the former, representing weak and strong epitopes respectively. Each peptide-position-allele occurrence is counted separately.

To weight low-rank peptides more heavily, the system calculated a cumulative immunogenicity score by summing the number of peptide-position allele occurrences with EL percent ranks strictly below thresholds of 0.25%, 0.50%, 0.75%, 1.00%, 1.25%, 1.50%, 1.75%, and 2.00%. Because each prediction is counted once for every threshold it falls under, lower-rank predictions contribute more to the score. For instance, a rank of 0.2 is counted 8 times whereas a rank of 1.8 would only be counted once.

Six optimization strategies were used to expedite searching without changing peptide count or the scoring function:

1. Once a parent sequence has been fully scored, only the peptides overlapping mutations were rescored for each candidate. Previously scored peptides were reused.
2. Repeated peptides within a scoring sequence were deduplicated and the NetMHCpan 4.1 model is only inference once.
3. Alleles are assessed in groups of 5 and the total peptide count was summed across all, candidates are rejected early once their accumulated peptide counts prove that they cannot satisfy the relative worsening to their parent requirement, described below.
4. Alleles with higher average counts in the parent sequence were assessed first to increase the effectiveness of early rejection.
5. Peptide-allele predictions were cached and reused throughout each search round.
6. The cache was retained between rounds so that predictions shared by related sequences could be reused.

#### 3. Fast Searching with MINT Cycling for Human Allele Deimmunization

During functional testing of the mouse alleles it was determined that all designs retained function when filtering (pLDDT ≥95 and RMSD ≤1 Å). The structural requirement was relaxed to pLDDT ≥90 and RMSD ≤2 Å for the more challenging redesign of human alleles, allowing broader exploration of sequence space.

Candidate sequences were assessed using the sum of the 8 threshold ranking buckets described above. The algorithm tracks two unique populations of protein sequences. First, is a list of 24 sequences representing the lowest immunogenicity hits (these are the running best discovered sequences through search, lowest immune score). In order to prevent the algorithm from getting trapped in a local minima and collapsing onto a set of very similar sequences, a second population of 24 is tracked attempting to maximize sequence diversity. The diversity set is rebuilt each round and contains twenty-four slots. Up to sixteen slots were populated using greedy selection of immune score with a sequence difference of at least two. The remaining eight slots were used to increase sequence diversity through min-max Hamming-distance selection. At each step, the minimum Hamming distance between each candidate and any already selected sequence was calculated, and the candidate with the largest minimum distance was added to the population. To prevent this diversity step from selecting poor designs, candidates were considered only if their immune score was no greater then, *S_best_* + *max*(10, ⌈0. 10 * *S_best_* ⌉) for deimmunization, where *S_best_* is the immune score of then best design found.

Before the search began, 512 ProteinMPNN sequences were generated from one initial backbone structure at a sampling temperature of 0.4 (see Design Library section of methods for more details on generating a backbone structure). The immune score was computed across all 512 redesigns and the best 96 were folded with AF3 and used along with the original sequence (97 designs in total) to seed the initial sequences contained within the previously mentioned lowest immunogenicity hits and diversity set of the search algorithm. In order to further introduce diversity during search, sixteen new global ProteinMPNN designs were generated every fifth round and used to update the lowest immunogenicity hits and diverse set to broaden sequence exploration, using the original backbone at temperature 0.4.

At each round of search, eight parent sequences were selected: the design with the best immune score and up to seven additional parents selected deterministically across the combined populations of lowest immunogenicity hits and the diverse set.

A hotspot was defined as a mutable position overlapped by at least one peptide with an EL percent rank below 2% for any tested allele. Hotspots were redesigned using three complementary strategies. First, exhaustive single substitutions were generated at each hotspot, excluding cysteine as a replacement. Second, to identify mutations that might work better together than individually, we selected up to eight pairs of mutable positions that co-occurred most frequently within predicted epitopes for each parent sequence. For each position, ProteinMPNN suggested up to three likely amino-acid substitutions. Substitutions with probabilities below 1% were excluded, and every combination of the remaining substitutions was tested across each pair (9 suggestions per pair and up to 72 suggestions across the 8 pairs). Finally, to introduce random mutations that push to unexplored sections of sequence space, joint ProteinMPNN resampling was run at temperature 0.4 on 2, 4 or 8 hotspot residues simultaneously, considering only the top 32 hotspots (residues overlapped by the most number of epitopes). These sampling strategies were repeated three, three and two times respectively, giving 8 joint samples total.

All of these mutation methods were applied to all parent sequences. This creates a pool of proposed next sequences. Each sequence is scored using the optimized search algorithm described earlier. To allow the search to reach improvements requiring multiple rounds of mutation, candidates were permitted to score slightly worse than their parents. Let (S_c) and (S_p) denote the child and parent scores, respectively. A candidate was made eligible for fold selection when its immune score was *S_c_* ≤ *S_p_* + *max*(2, ⌈0. 02 * *S_p_*⌉). Up to 36 best immune scoring sequences differing by at least 2 residues were selected for folding. Then the aforementioned min-max hamming distance algorithm was used to select up to 48 designs for folding.

Of the eligible candidates, up to 48 were then folded using AF3. Search folds used five recycles and one diffusion sample, with empty paired and unpaired MSAs and no templating. A candidate is retained if it meets the structural requirement of Cα pLDDT ≥90 and aligned Cα RMSD ≤2.0 from the original structure. The designs passing the structural filter are entered into the lowest immunogenicity hits and diversity set and this process is repeated.

## 4. Termination & Confirmation

Search can be run in parallel across multiple backbones and sequence starting points to explore differing regions of sequence space. To deimmunize Luxsit-i 20 such lineages were run in parallel. The top candidate from each lineage underwent independent peptide rescoring and a separate AF3 confirmation folds to assess whether each met the required structural cutoff. Search could also terminate early if a zero deimmunization score was independently confirmed and all additional AF3 folds passed the structural criteria.

### Sequence entropy analysis

To explore the sequence diversity of the Luxsit-I protein structure, MPNN was used to propose 10,000 redesigned Luxsit-I sequences at a temperature of 0.4. Following structure prediction with AF3 321 designs met the structure criteria (mean Cα pLDDT ≥ 90 and aligned Cα RMSD ≤ 2.0 Å). Sequence diversity was quantified using Shannon entropy across each of the 117 residue positions (Figure 1SA).

### Benchmarking

Both the initial non-cycling method and the cycling method were benchmarked against the same panel of 25 HLA class-I alleles. A starting pool of 20 candidates was selected from the pool of confident designs (Design Library) and optimized with both methods. The iterative method was allowed to run for 5 rounds across all 20 designs. The cycling method treated each of the 20 designs as an independent population and ran each for up to 320 rounds of search. Both used the same structural acceptance criteria (mean Cα pLDDT ≥ 90, aligned Cα RMSD ≤ 2.0 Å). For each method, cumulative cpu hours and the best redesigned candidates were recorded. Because per-round cost differs between the two methods, CPU hours, and not rounds, is the matched axis for comparison (Figure 1F).

### Ablation Analysis

To determine the speed of peptide searching using the six optimizations described above, a dataset of 600 searched sequences was collected from 3 rounds of cycling using MINT. Then, completely unoptimized, cached search and fully optimized versions were tested and the number of inferences of NetMHCpan was evaluated to determine the effect of each method on the resulting netMHCpan4.1 search counts.

---

**Figure S1.**
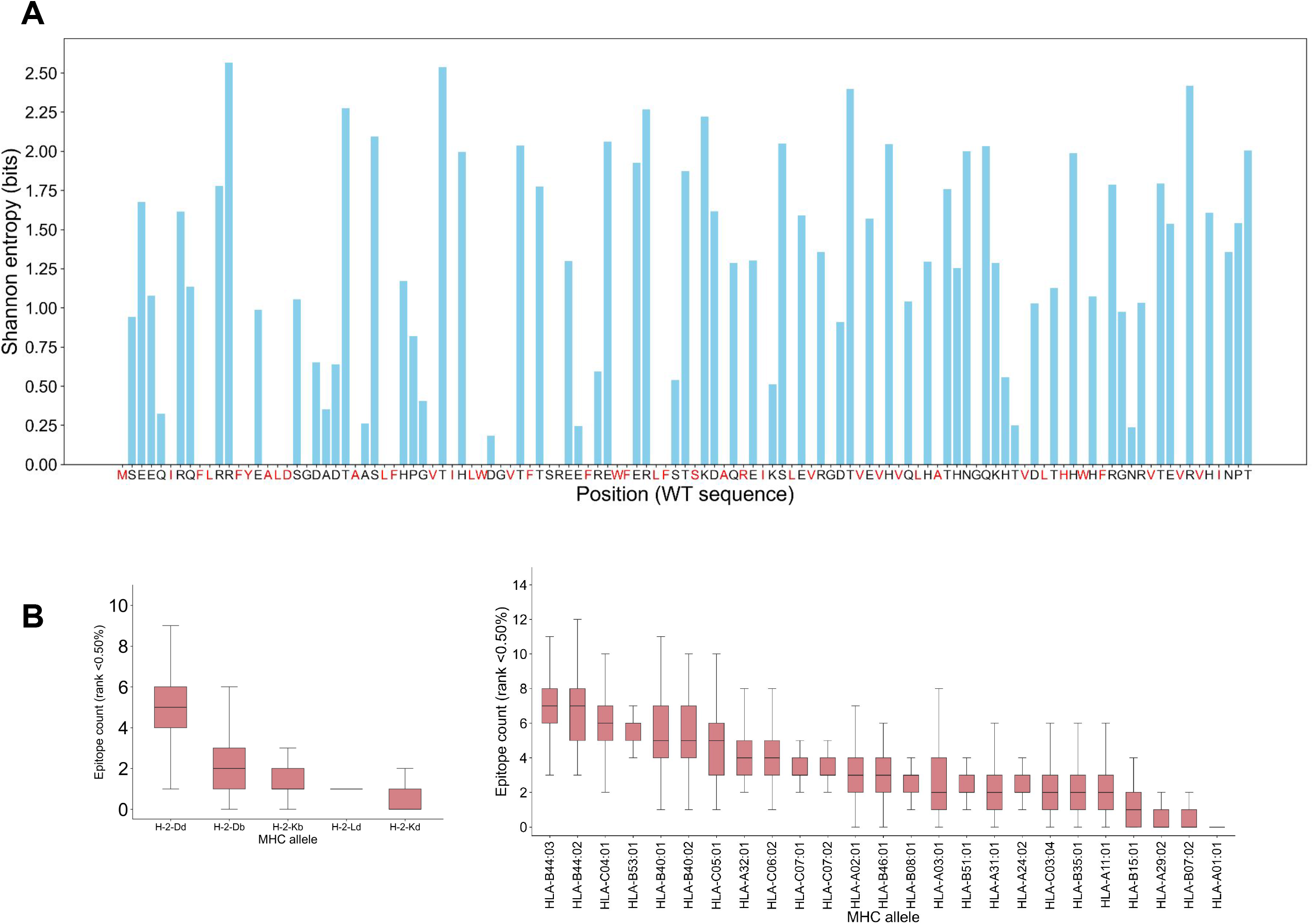
Sequence diversity and predicted MHC-I epitope burden of LuxSit-i variants. (A) Sequence entropy of each site illustrates amino acid diversity at each position among LuxSit-i sequences generated from MINT with no cycling. Red color indicating the fixed residues. (B) Distribution of the number of strong (EL rank <0.5%) MHC-I epitopes for each of 5 mouse MHC-Is (left) and each of 25 human MHC-Is (right) among LuxSit-i sequences generated from MINT with no cycling.

**Figure S2.**
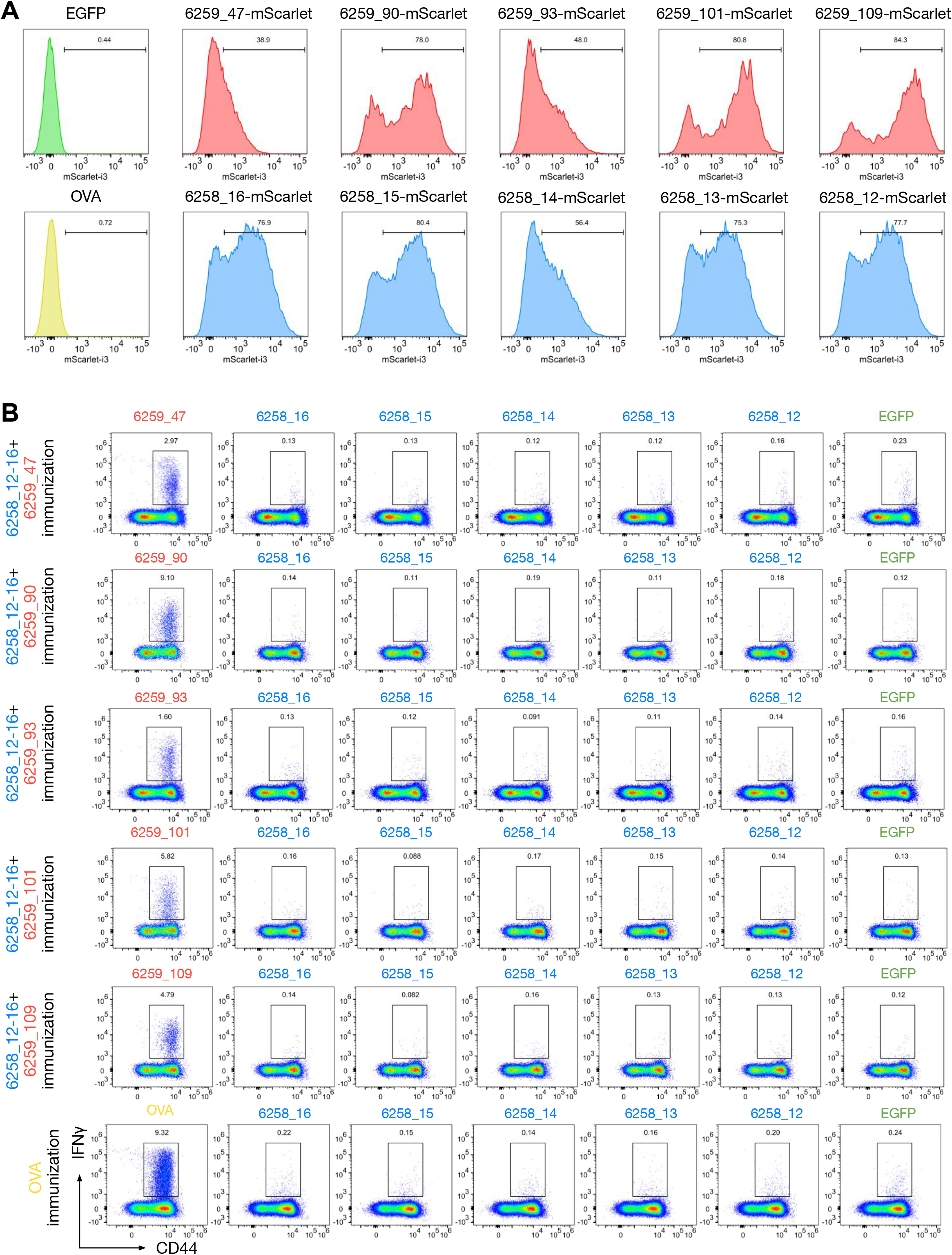
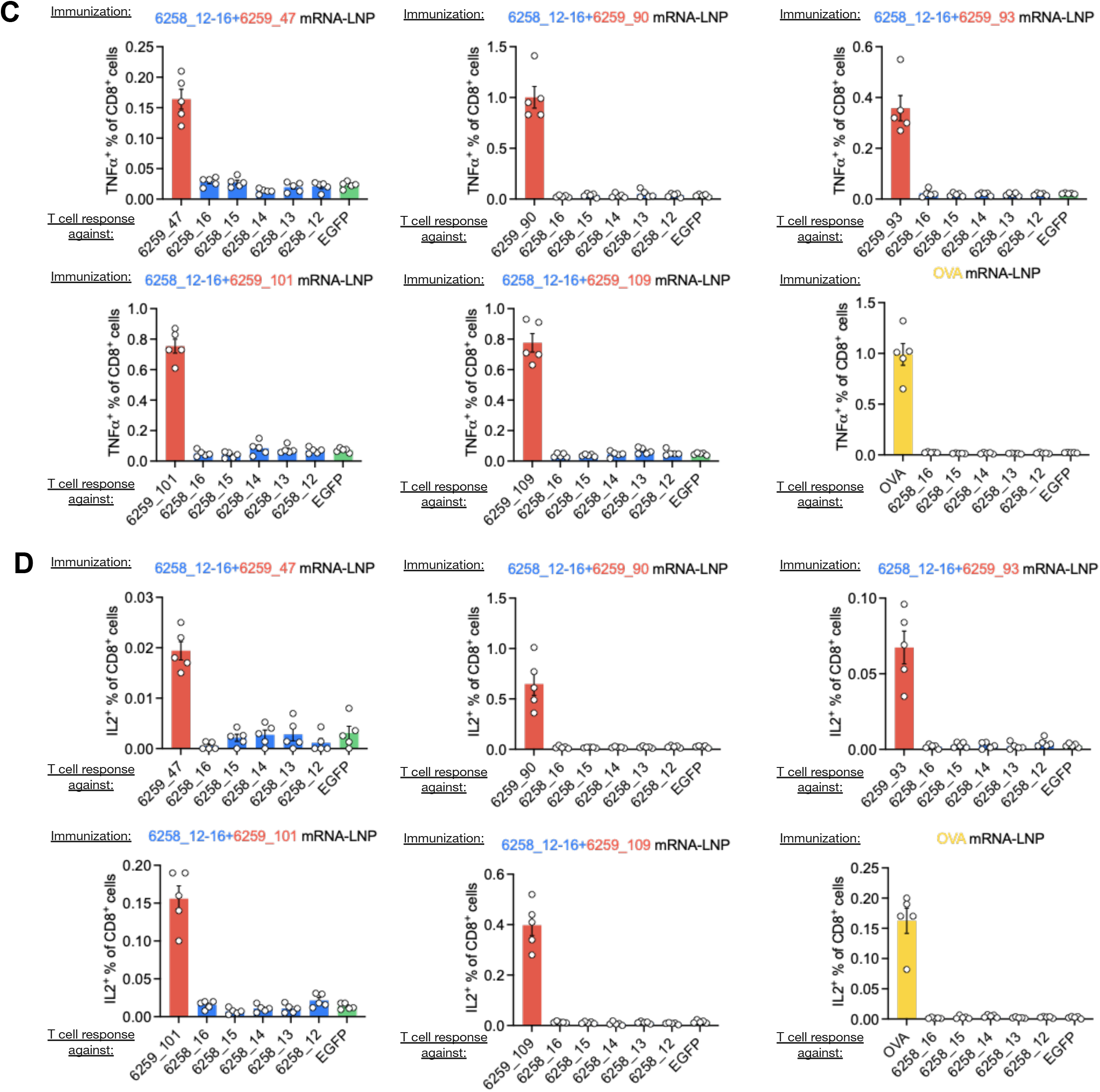
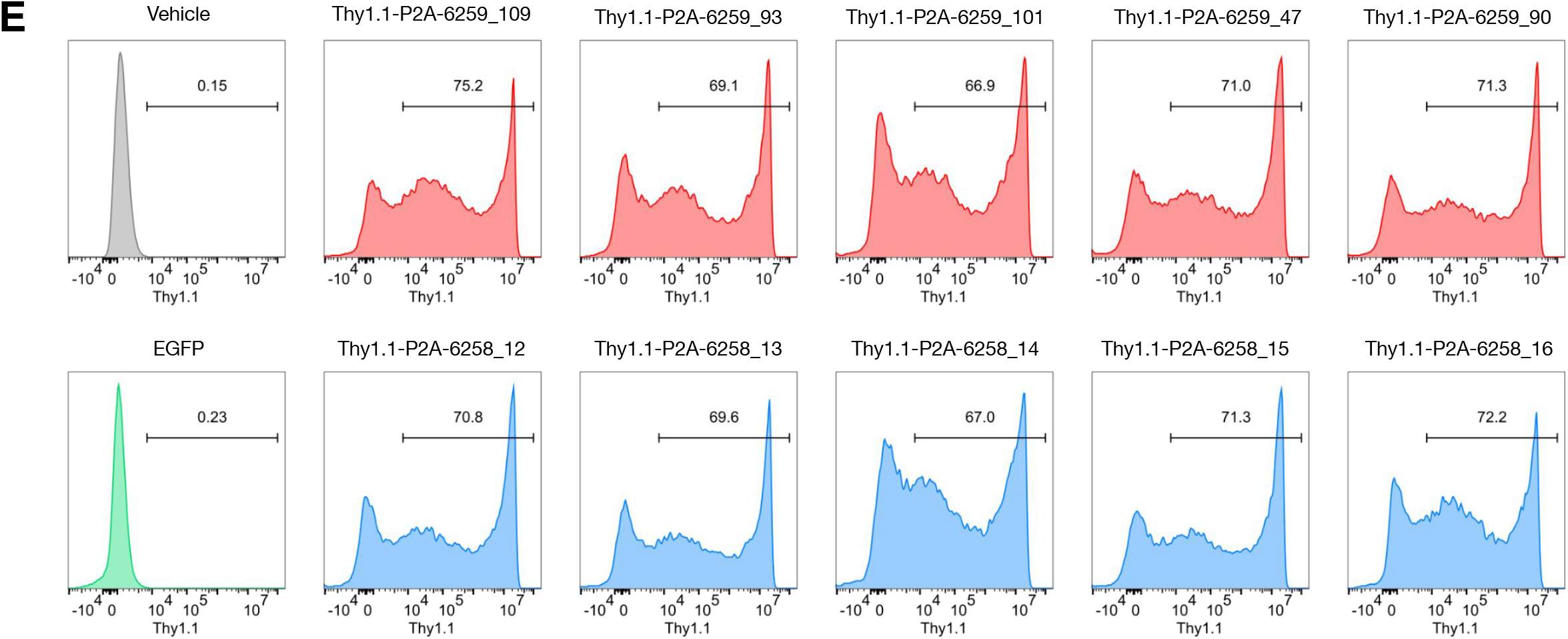
Expression and characterization of CD8⁺ T cell responses to designed luciferase variants. (A) mScarlet reporter expression in bone marrow-derived dendritic cells (BMDCs) transfected with mRNAs encoding individual LuxSit-i variants. (B) Representative flow cytometry plots showing IFN-γ production by CD8⁺ T cells following immunization with 5+1 fusion constructs. (C) Frequencies of TNF-α⁺ CD8⁺ T cells responding to individual LuxSit-i variants following 5+1 fusion immunization. (D) Frequencies of IL-2⁺ CD8⁺ T cells responding to individual LuxSit-i variants following 5+1 fusion immunization. (E) Thy1.1 reporter expression in bone marrow-derived dendritic cells (BMDCs) transfected with mRNAs encoding individual LuxSit-i variants.

**Table S1.**
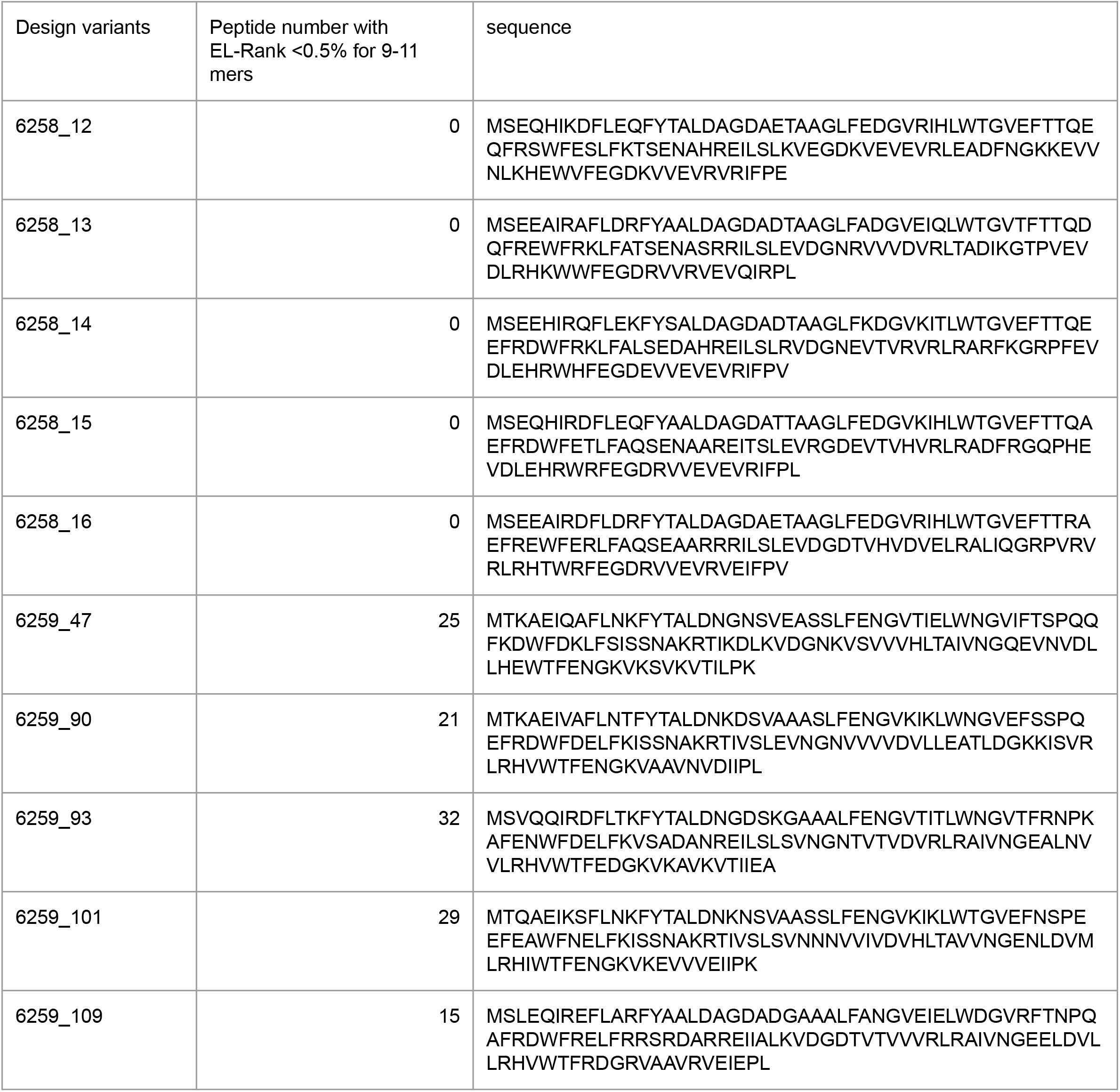
Sequences and numbers of predicted immunogenic peptides for the 10 LuxSit-i variants used in the in vivo experiments.

